# Computational Binding Affinities of Disheveled PDZ Protein-Ligand Complexes

**DOI:** 10.1101/2025.09.21.672002

**Authors:** Aarav Singh, Harsha Kancharla, Andrew Jubintoro, Patrick Blankenberg, Jie Zheng

**Affiliations:** Stein Eye Institute, Department of Ophthalmology, David Geffen School of Medicine at UCLA, Los Angeles, CA 90095, USA; Department of Integrative Biology and Physiology, UCLA, Los Angeles, CA 90095, USA

## Abstract

Wnt/ß-catenin signaling is critical for cell growth and development, with its hyperactive dysregulation implicated in the development of cancer. Current therapeutic research on inhibition of Wnt/ß-catenin signaling is impeded by the high cost of experimentally determining binding affinities. Consequently, interest has risen in screening potential inhibitors binding affinities with computational tools to reduce costs. Here, we test the validity of a computational molecular dynamics simulator, Binding Free Energy Estimator 2 (BFEE2), for determining peptide ligand affinity for Wnt/ß-catenin signaling. We focus on the Dishevelled (DVL) PDZ domain, a key mediator in WNT signaling through its ability to bind to various peptide ligands. We analyze the binding affinities of several DVL PDZ domain-peptide against previously established results to determine the validity of computational analysis. We conclude that computational molecular dynamics simulations were successful for peptide-ligand complexes.

## 1. INTRODUCTION

The Wnt signaling pathway is highly conserved across multicellular animals and is crucial in regulating cell proliferation, cell fate decisions, and embryonic development. In the canonical cascade, Wnt ligands bind Frizzled receptors, which recruit Dishevelled-1 (DVL1). DVL1 is an intracellular signaling protein that gets recruited and helps carry the signal forward; the protein also subsequently promotes B-catenin,a protein that helps cells adhere in Wnt signalling and Wnt-dependent gene activation stabilization [^1^].

However, overactivation of DVL1 can drive excessive B-catenin accumulation and uncontrolled cell proliferation. This hyperproliferation contributes to tumorigenesis, in particular retinoblastomas. This disease accounts for about 3% of cancers in children younger than 15 years, with an age-adjusted annual incidence of 18.4 cases per million in children aged 0–4 years in the United States[^1^]. Identifying potent peptide inhibitors that target the DV1 PDZ domain is critical in restoring normal Wnt signaling and cellular growth [^2,3^]. An accurate pre-screening tool would enable more efficient evaluation of new ligands, helping to prioritize those with stronger binding affinities to the Dishevelled protein. Existing pre-screening tools such as molecular docking can rapidly rank candidate ligands with much lower computational cost than molecular-dynamics based simulations; however, because docking scores functions that do not fully capture the thermodynamic complexity of binding, they are therefore less reliable for accurate quantitative binding affinity prediction.

Here we introduce Binding Free Energy Estimator 2 (BFEE2), a protein-ligand binding affinity tool built on nano-scale molecular dynamics (NAMD) simulations that estimates the free energy of binding between a ligand and its protein. Binding Free Energy Estimator 2 (BFEE2) was tested for validity as a pre-screening tool for the DVL domain. BFEE2 uses the geometric-restraint route via fixing the ligand’s orientation with distance, angle, and dihedral restraints, samples the restrained complex while gradually decoupling the ligand. The restraints are then analytically released to yield the standard-state binding free energy [^4^]. Twenty-two DVL protein-ligand structures were initially considered for analysis. Following screening, four peptide-ligand DVL complexes met the inclusion criteria and were selected for full analysis. Complex structure files were sourced from the RCSB Protein Data Bank and parameterized according to standard NAMD preparation procedures. The parameterized peptide-ligand complexes were then analyzed using BFEE2 to generate computational binding affinity estimates. These in silico binding affinities were compared with published experimental binding affinity values to assess agreement between computational and experimental results. The peptide-ligand complexes showed convergence between experimental and computational binding affinities, supporting BFEE2 as a useful computational tool for estimating peptide-ligand binding affinities in DVL systems.

## 2. METHODS

### 2.1 Inclusion Criteria

DVL domain complexes were sourced from RCSB Protein Data Bank [^5^]. DVL domain complexes which lacked ligands were excluded for analysis given BFEE2 is a tool for modeling ligand affinity interactions. DVL domain complexes which had incomplete structure or no peer-reviewed experimental binding affinities were excluded from analysis.

### 2.2 Pre-Processing

Artifact water and solvent ions were pruned from complex protein database files to reduce X-Ray/NMR noise. Dimerized complexes were converted into monomers by analyzing the protein from one monomer and a ligand from the other monomer to create a single protein-ligand complex.

Atomic structure topology was generated by standard NAMD analysis. Molecular topology files from NAMD were used to construct an atomic connectivity file in a .psf format for analysis with Visual Molecular (VMD) AutoPSF. The complex .pdb and .psf molecule files were padded with 10 Angstroms to prevent interactions with periodic images and allow for realistic solvation. The system was solvated to 0.15 M NaCl to mimic biologic conditions [^6,7^].

### 2.3 BFEE2 Analysis

The CHARMM36 peptide parameter file was obtained from NAMD for the peptide complexes.

As seen in Figure 1, each step of the BFEE2 protocol targets a specific component of the ligand’s binding affinity by applying Colvars, a software library used in MD simulations to bias runs, to sample the energy landscape. Three different forms of restraints were used: harmonic, angular, and RMSD. A harmonic distance restraint keeps the ligand near the binding pocket by constraining the center-of-mass distance between the protein and ligand. Angular restraints limit the ligand’s rotational freedom relative to the protein, enabling systematic sampling of orientational degrees of freedom such as eulerTheta, eulerPhi, eulerPsi, polarTheta, and polarPhi. The RMSD restraint ensures the ligand remains within a defined structural envelope, distinguishing bound from unbound states [^4,8^].

**Figure 1.**
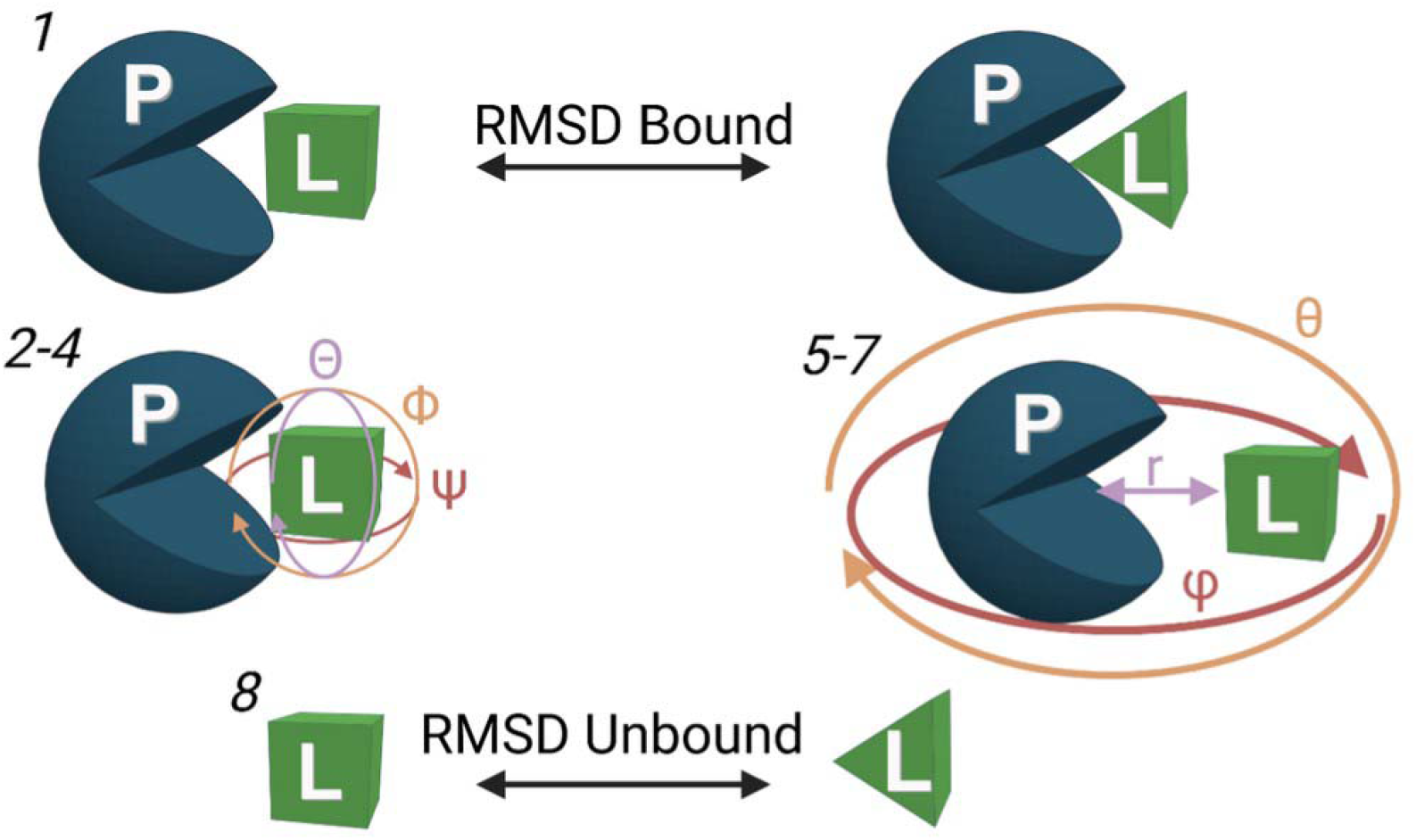
BFEE2 geometric route setup adapted from BFEE2 paper; steps 1-8 along with their respective restraints [^11^].

As part of Colvars, BFEE2 used the Adaptive Biasing Force (ABF) method to provide enhanced sampling. ABF identifies which degrees of freedom, like ligand rotation or separation, are energetically unfavorable then applies a small, adaptive biasing force to help the molecule overcome these energy barriers and explore the configurational space better.

As seen in Figure 2, after running 5,000,000 steps in a BFEE2, a potential of mean force (PMF) is generated, displaying how the free energy changes along a selected collective variable (e.g. distance, angle, or orientation between protein and ligand). The binding free energy is then calculated using Boltzmann-weighted integration, which evaluates the probability of the ligand being bound versus unbound to yield the absolute binding free energy. Each step of the BFEE2 simulation corresponds to the free energy associated with a specific restraint, and summing up these values gives the total binding free energy of the ligand to the protein. To ensure the simulation ran properly, two key checkpoints were used. First, in the 000_eq (equilibration) folder, the DCD trajectory file was loaded into VMD alongside the corresponding PSF structure file to visually confirm that the ligand remained in the binding pocket and did not drift or unbind prematurely, which is a sign of an equilibrated system. Second, for each simulation window, the RMSD plot was inspected to verify convergence, which is indicated by RMSD values flattening out gradually towards run completion (Figure 2).

**Figure 2.**
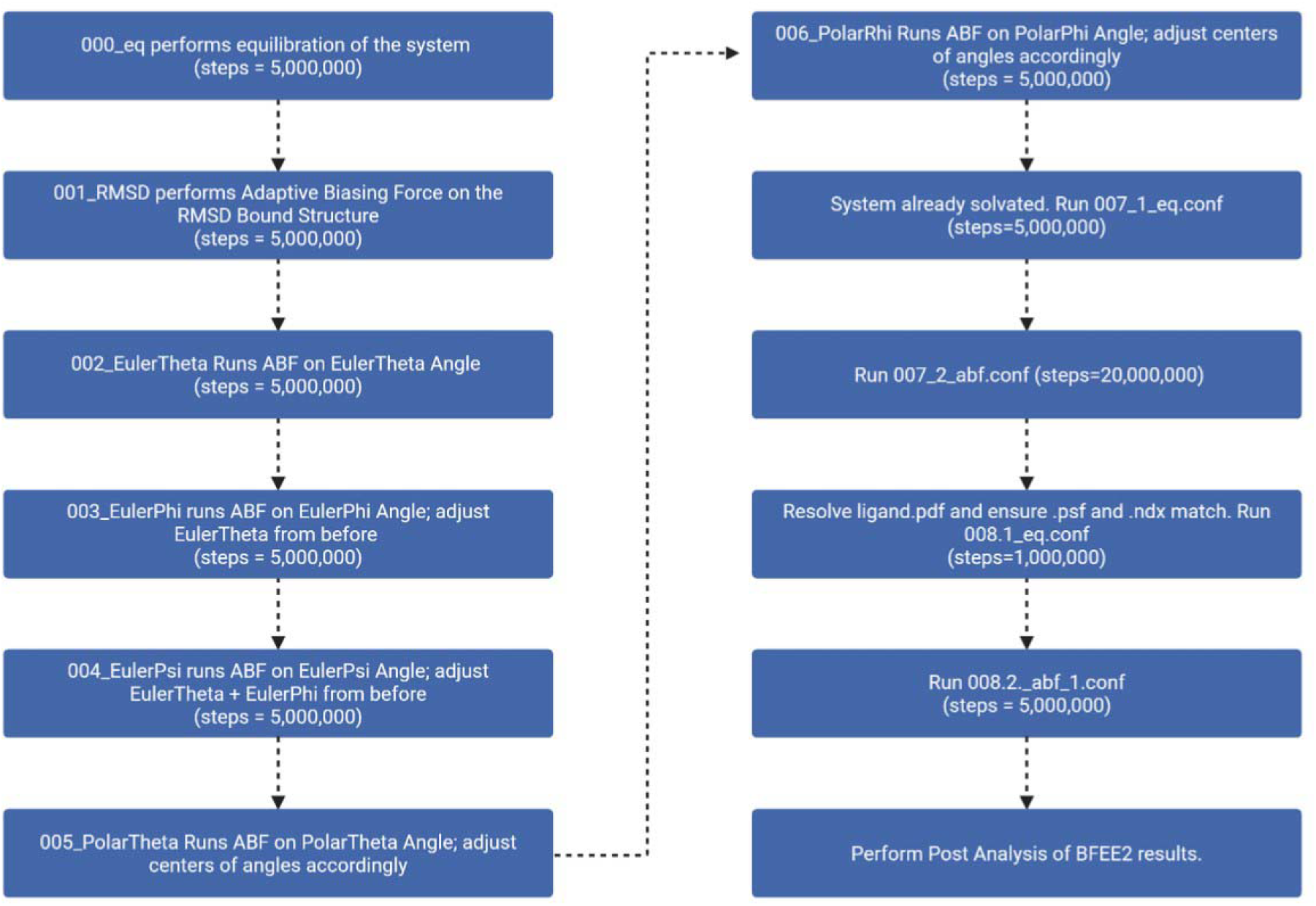
The default parameters of each step of the BFEE2 simulation are outputted with their corresponding steps. For small protein ligand complexes, the default parameterization approximately reaches RMSD convergence.

Computational binding affinities were compared to experimentally obtained values obtained from the published data on RCSB. Differences in binding affinities less than 1 kcal/mol were considered to be convergent while differences outside this range were considered to be divergent [^4^].

## 3. Results

22 DVL protein structures were initially screened for inclusion in the analysis. Structures were excluded if they lacked a bound ligand complex, had incomplete publicly available structural information, or did not have verified experimentally determined binding affinity data. Following this screening process, four peptide-ligand DVL complexes met the inclusion criteria and were selected for full BFEE2 analysis: 1L6O, 3CC0, 3CBY, and 3CBX.

Protein-peptide complex binding affinities calculated in silico were found to be congruent with verified experimental values, within ±1 kcal/mol. This indicates that the BFEE2 simulations successfully converged for peptide-ligand complexes.

## 4. Discussion

As seen from Table 1, BFEE2 binding affinity simulations were found to be relatively accurate when applied to peptide-PDZ complexes. This simulation validates the BFEE2 protocol for peptide ligand complexes. This accurate parameterization allows simulation to generate experimental measurements when only peptide-PDZ complexes are considered. Analysis of peptides with BFEE2 is likely accurate across different domains due to peptides simply being a collection of amino acids. Amino acid forcefields have been rigorously studied, and consequently BFEE2 simulations conducted with peptides are the most promising.

**Table 1.**
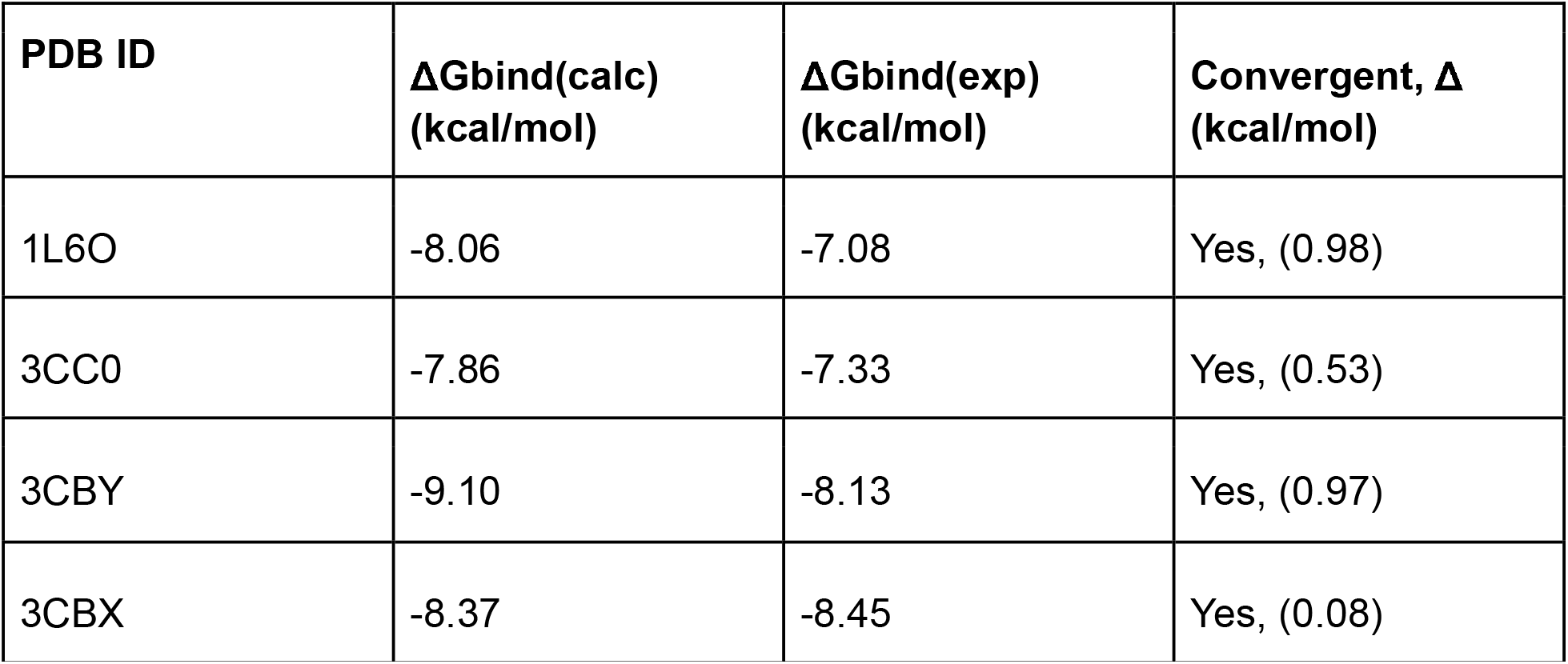
Comparison of computational vs experimental binding affinities for all protein-ligand complexes [^2^].

## 5. Conclusion

We conclude that BFEE2 is a useful pre-screening tool for evaluating dishevelled peptide ligands, with the four analyzed peptide-ligand complexes successfully converging and showing strong agreement with experimentally determined binding affinity data. These results suggest that the BFEE2 geometric route can provide a practical computational workflow for estimating peptide binding affinities in DVL PDZ systems.

The optimized BFEE2 workflow did not require extensive manual parameterization for the peptide systems analyzed, making it accessible for researchers studying dishevelled peptide ligands or similar peptide-based protein-ligand interactions. However, convenient use of this workflow still requires access to a strong GPU or supercomputing cluster because the required NAMD simulations are computationally intensive.

More broadly, these findings support the use of BFEE2 as a peptide-ligand screening approach, particularly when experimental binding data are limited or when prioritizing candidate ligands for future study. Future work should expand this analysis to a larger set of peptide complexes and compare the geometric route with BFEE2’s alchemical option to determine whether similar or improved agreement with experimental binding affinities can be achieved.

## SUPPLEMENTARY MATERIALS

Below outlines the issues encountered with each protein-ligand complex. This is included for future work on the peptides and small molecule ligand complexes particularly.

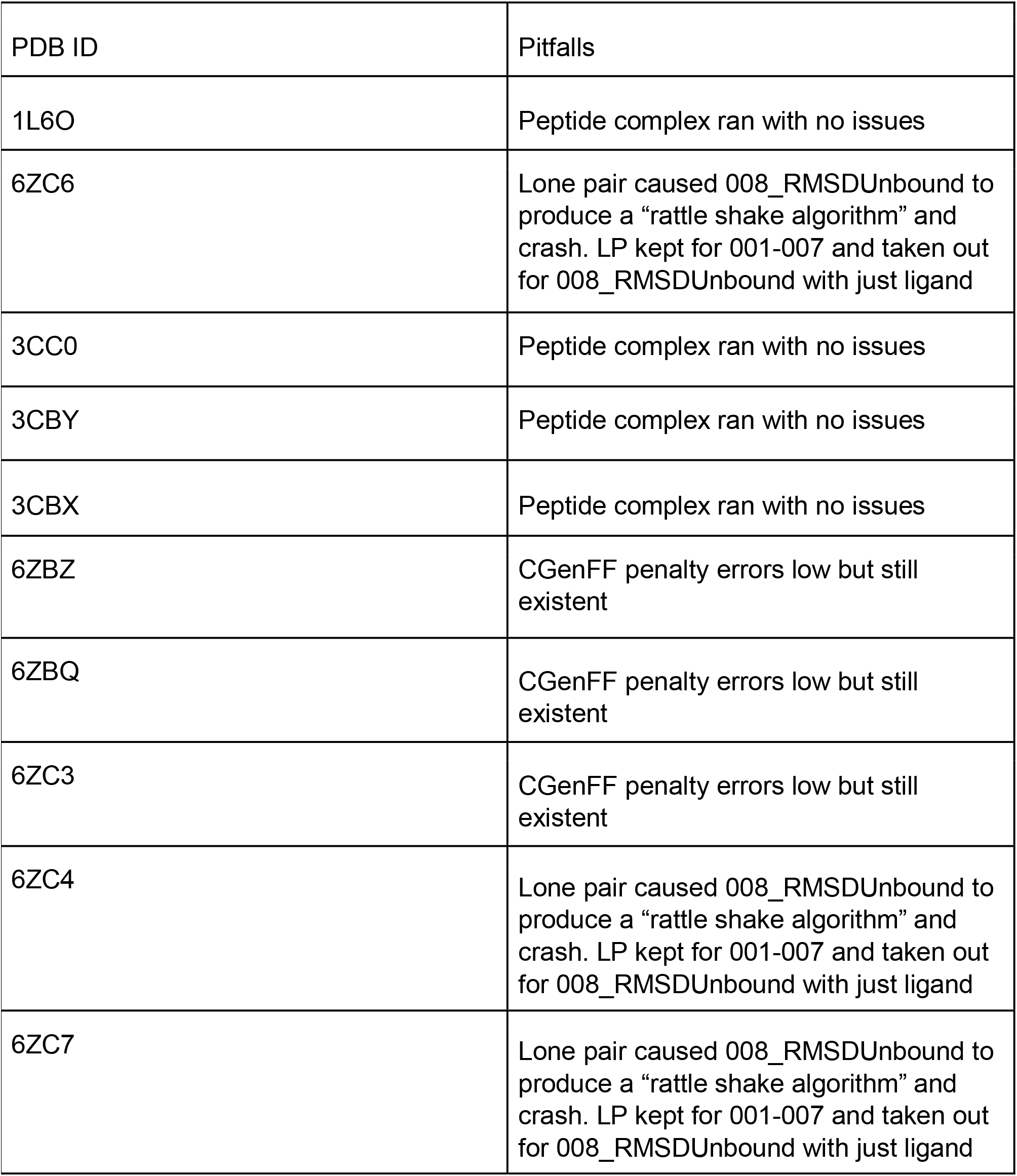
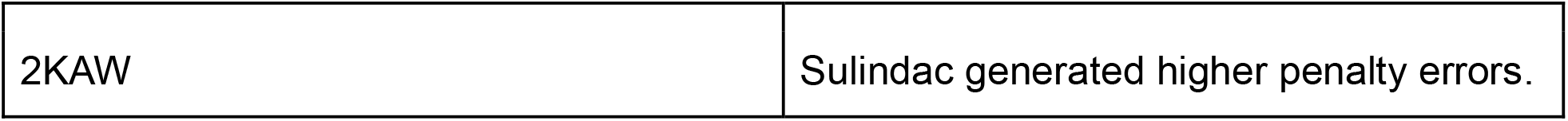

